# The reducing end of cell wall oligosaccharides is critical for DAMP activity in *Arabidopsis thaliana* and can be exploited by oligosaccharide oxidases in the scavenging of phenolic radicals

**DOI:** 10.1101/2024.07.30.605788

**Authors:** Moira Giovannoni, Anna Scortica, Valentina Scafati, Emilia Piccirilli, Daniela Sorio, Manuel Benedetti, Benedetta Mattei

## Abstract

- The enzymatic hydrolysis of cell wall polysaccharides results in the production of oligosaccharides with nature of damage-associated molecular patterns (DAMPs) that are perceived by plants as danger signals. *In vitro*, the oxidation of oligogalacturonides and cellodextrins by plant FAD-dependent oligosaccharide-oxidases (OSOXs) suppresses their elicitor activity, suggesting a protective role of OSOXs against a prolonged activation of defense responses potentially deleterious for plant health. However, OSOXs are also produced by phytopathogens and saprotrophs, complicating the understanding of their role in plant-microbe interactions.
- The oxidation catalyzed by specific fungal OSOXs also converts the elicitor-active cello-tetraose and xylo-tetraose into elicitor-inactive forms, indicating that the oxidation state of cell wall oligosaccharides is crucial for their DAMP function, irrespective of whether the OSOX originates from fungi or plants.
- Certain OSOXs can transfer the electrons from the reducing end of these oligosaccharides to phenolic radicals instead of molecular O_2_, highlighting an unexpected sub-functionalization of these enzymes.
- The activity of OSOXs may be crucial for a thorough understanding of cell wall metabolism since these enzymes can redirect the reducing power from sugars to phenolic components of the plant cell wall, an insight with relevant implications for both plant physiology and green technologies.

## Introduction

Plant cell wall is an extracellular structure composed of polysaccharides, proteins and phenolic compounds. It plays a fundamental role in physiological processes, including the maintenance of turgor pressure, cell shape, and protection against microbes (Scortica et al., 2022). In addition, plant cell wall can also function as a reservoir of energy, with its reducing power locked within its complex architecture and heterogenous composition. To colonize the plant tissue, phytopathogens need first to break down the different components of the cell wall, whose degradation is achieved through the secretion of cell wall-degrading enzymes, a class of enzymes mainly constituted by glycosyl hydrolases, oxidoreductases and lyases (Giovannoni et al., 2020). The enzymatic hydrolysis of cell wall polysaccharides results in a transient accumulation of cell wall fragments with nature of damage-associated molecular patterns (DAMPs) such as oligogalacturonides (OGs) and cellodextrins (CDs), that plants perceive as danger signals (Pontiggia et al., 2020). These cell wall oligosaccharides may also act as negative regulators of various developmental processes. Their activity is crucial for balancing the growth-defense trade-off due to their contrasting effects on immunity and growth (Pontiggia et al., 2020; Scortica et al., 2022). It is important to bear in mind that, although DAMP-triggered immunity confers protection against invading microbes, an exaggerated and prolonged activation of defense negatively affects plant health. A clear effect of this process can be observed in the stunted growth-phenotype of *Arabidopsis thaliana* plants that accumulate OGs in uncontrolled manner (Benedetti et al., 2015). *In vitro*, the oxidation of OGs and CDs by plant FAD-dependent oligosaccharide-oxidases (OSOXs) strongly reduces their elicitor activity, suggesting a protective role of OSOXs against a self-deleterious hyper-activation of DAMP-triggered immunity (Benedetti et al., 2018; Costantini et al., 2024; Locci et al., 2019). In accordance with this hypothesis, Arabidopsis plants treated with oxidized OGs and CDs showed a lower defense gene induction, a weaker oxidative burst, and a reduced accumulation of callose compared to plants treated with the corresponding non-oxidized elicitors (Benedetti et al., 2018; Costantini et al., 2024; Locci et al., 2019). OSOXs belong to the family of berberine bridge enzyme-like (BBE-l) proteins, a class of flavoenzymes widely distributed in bacteria, fungi and plants (Daniel et al., 2017). Different OSOXs from *A. thaliana* have been identified as specific oligogalacturonide-oxidases (OGOXs) and cellodextrin-oxidases (CELLOXs) (Benedetti et al., 2018; Costantini et al., 2024; Locci et al., 2019). OGOXs include four isoforms (OGOX1-4) that oxidize OGs, whereas CELLOXs include two isoforms (CELLOX1-2) that oxidize CDs. Interestingly, OSOXs are also produced by phytopathogens and saprotrophs such as the gluco-oligosaccharide oxidase from *Sarocladium strictum* (GOOX) (Lee et al., 2005), active on CDs as plant CELLOXs, and the xylo-oligosaccharide oxidase from *Miceliophtora thermophila* (XOOX), active on xylo-oligosaccharides (Ferrari et al., 2016). Thus, the existence of OSOXs in both fungi and plants complicates the comprehension of their role in plant-microbe interaction. In the last 20 years, the different OSOXs so far identified have been described as specific oxidases, i.e., oxidoreductases capable of transferring electrons from the reducing end of oligosaccharides to molecular O_2_, thereby producing H_2_O_2_. According to the stoichiometry of enzymatic reaction, one molecule of H_2_O_2_ is produced for each C1-oxidized oligosaccharide. However, in the presence of short oligosaccharides and specific electron acceptors, both OGOX1 and CELLOX1 also act as conditional dehydrogenase, for example, by transferring the electrons from the reducing end of tetra-galacturonic acid (OG4) and cello-tetraose (CD4), respectively, to the radical cation ABTS^•+^ (single-electron transfer = dehydrogenase activity) rather than to molecular O_2_ (two-electron transfer to O_2_ = oxidase activity) (Scortica et al., 2023).

Recently, 3-α-L-arabinofuranosyl-xylo-tetraose (XA3XX) and xylo-tetraose (Xyl4) have been reported to act as DAMPs in *A. thaliana* (Fernández-Calvo et al., 2024; Mélida et al., 2020). By using OSOXs of both fungal and plant origin, we demonstrate here that oxidized Xyl4 (xylo-tetraonic acid, ox-Xyl4) and oxidized CD4 (cello-tetraonic acid, ox-CD4) have a weaker elicitor activity than their non-oxidized counterparts, reinforcing the view that DAMP action of cell wall oligosaccharides is dependent on the oxidation state of their reducing end, regardless whether the OSOX employed for their oxidation originates from fungi or plants. In addition, we also prove that certain OSOXs redirect the electrons from the reducing end of cell wall oligosaccharides to oligo-guaiacol radicals, instead of molecular O_2_. This sub-functionalization, based on the preference for different electron acceptor substrates (e.g., O_2_ *vs* oligo-guaiacol radicals), may explain the presence of different OSOXs with similar substrate specificities (i.e., capability of subtracting electrons from the same oligosaccharide), generally referred to as OSOX paralogs, in the same organism (Benedetti et al., 2018; Costantini et al., 2024)

## Materials and Methods

### Design of the constructs expressing plant and fungal OSOXs in *Pichia pastoris*

The sumoylated-flag-his tagged form of cellodextrin-oxidase 1 from *Arabidopsis thaliana* (FHS-CELLOX1) was cloned in pPICZαB according to the same strategy described in (Costantini et al., 2024; Scortica et al., 2022). The gene encoding the fungal OSOXs i.e., the xylo-oligosaccharide oxidase (XOOX) from *Myceliophthora thermophila* C1 and the gluco-oligosaccharide oxidase (GOOX) from *Sarocladium strictum* were codon-optimized according to the codon usage of *Komagataella phaffii*, formerly known as *Pichia pastoris*, using the software OPTIMZER (http://genomes.urv.es/OPTIMIZER/) (Puigbò et al., 2007). Putative signal peptides of both fungal OSOXs were identified by using the SignalP-5.0 server (http://www.cbs.dtu.dk/services/SignalP/) (Almagro Armenteros et al., 2019) and excluded from the sequences. The extra-bases encoding the restriction sites of PstI and XbaI were added at the 5^I^ and 3^I^ ends of such sequence and used to clone the two synthetic genes in pPICZαB (Invitrogen, San Diego, CA) in frame with the sequence encoding the 6xHis-tags (HHHHHH) and the yeast α factor for secretion in the culture medium. The entire sequences were synthesized by Genescript (https://www.genscript.com/). The tagged version of XOOX and GOOX here described will be referred to as H-XOOX and H-GOOX throughout the manuscript.

### Preparation of oxidized Xyl4 (ox-Xyl4) and oxidized CD4 (ox-CD4)

1 mg of commercial Xyl4 (O-XTE, Megazyme) and CD4 (O-CTE, Megazyme) was separately dissolved in 0.5 mL of 20 mM Tris-HCl pH 7.0 and 20 mM NaCl. The mixtures were incubated at 30 ⁰C for 16 h by adding the purified OSOXs (0.04 g.L^-1^ each), i.e., H-XOOX for Xyl4, H-GOOX and FHS-CELLOX1 for CD4, and catalase (0.02 g.L^-1^) from bovine liver (Sigma). After incubation, a small aliquot of each reaction was analyzed by HPLC in order to verify the oxidation state of Xyl4 and CD4. HPLC analysis was performed with the same instrumentations and conditions described in (Scafati et al., 2022) by modifying the amount of sugar standards used in the analysis (2 mg.mL^-1^) and the volume injection of each sample (5 µL). All the data acquired were processed by Shimadzu LabSolutions control software. The final reactions were loaded into a Vivaspin® 500 (10000 MWCO PES) and then centrifuged (5 min, 4000 ×g) to separate OSOXs and catalase from the oxidized sugars. The filtrated sugars were quantified by using total sugar assay (Dubois et al., 1956) and the reducing sugar assay (Lever, 1972). As controls, the same procedures were carried out on 1 mg of Xyl4 and CD4 by adding heat inactivated OSOXs and catalase.

### MAPKs immunoblot analysis

Ten day-old *A. thaliana* seedlings were treated with different oligosaccharide-based elicitors (80 μg.mL^-1^ each), i.e., Xyl4, ox-Xyl4 (H-XOOX), CD4, ox-CD4 (H-GOOX), ox-CD4 (FHS-CELLOX1), and with flg22 (10 nM) as positive control, for 5 and 10 minutes respectively. 20 seedlings (about 100-200 mg FW) were pulverized in liquid nitrogen and the total proteins were extracted with 100 µl of phosphatase-inhibiting buffer for 20 min (4°C) and then centrifuged at 13000 x g for 30 min (4°C), in accordance with (Giovannoni, Marti, et al., 2021). The supernatant was recovered, and the protein extracts (40 μg for each sample) were separated using 10% SDS-PAGE. For detection of phosphorylated MAPKs an anti-p44/42-ERK antibody was used (1:2000, Cell Signaling Technology). For detection of total MAPKs, the anti-AtMPK3 (1:2500, Sigma, M8318) and anti-AtMPK6 (1:10000, Sigma, A7104) antibodies were used. Nitrocellulose membranes were then incubated in Tris-buffered saline, 0.1% Tween 20 and 0.5% (w/v) BSA with a secondary horseradish peroxidase-conjugated anti-rabbit antibody (1:6000, Bio-Rad). The chemiluminescence-based detection was performed by using the Clarity™ Western ECL substrate detection kit (Bio-Rad) in accordance with (Giovannoni, Lironi, et al., 2021).

### RNA extraction and gene expression analysis

10-day-old seedlings were treated with each oligosaccharide-based elicitor (80 µg.mL^-1^), i.e., [Xyl4, ox-Xyl4 (H-XOOX), CD4, ox-CD4 (H-GOOX), ox-CD4 (FHS-CELLOX1), and flg22 (10 nM) for 1 h. Plant tissues were grinded with Retsch MM 301 vibratory mill for 1 min at 24 Hz. NucleoZOL one phase RNA purification reagent was used to extract the total RNA from samples following the manufacturer’s specification. The cDNA synthesis and quantitative real-time PCR (qRT-PCR) analyses were performed in accordance with (Giovannoni, Lironi, et al., 2021). The gene expression analysis was performed on three different biological replicates (each composed of three technical replicates, n=3) mediating the results obtained. The primers used for qRT-PCR analysis are listed in Table S1.

## Results

### Heterologous expression of plant and fungal OSOXs and preparation of oxidized tetra-saccharides

We expressed the flag-his-Sumo-start tagged version of CELLOX1 from *A. thaliana* (FHS-CELLOX1), the his-tagged version of GOOX from *S. strictum* (H-GOOX) and the his-tagged version of XOOX from *M. thermophila* (H-XOOX) in *Komagataella phaffii* – formerly known as *P. pastoris* (Heistinger et al., 2020). FHS-CELLOX1 and H-GOOX are both active on CDs whereas H-XOOX is active on xylo-oligosaccharides (Ferrari et al., 2016; Lee et al., 2005; Locci et al., 2019). FHS-CELLOX1 and H-XOOX were purified from the culture filtrate of recombinant yeasts by using a 6xhis-tag affinity chromatography. As already reported (Lee et al., 2005), the Ni-column was not able to bind H-GOOX, therefore its purification was carried out by using a hydrophobic interaction chromatography (HIC). The quality of purified FHS-CELLOX1, H-XOOX and H-GOOX was assessed by SDS-PAGE and immuno-decoration analyses (Fig. **1a**). Subsequently, the purified FHS-CELLOX1 and H-GOOX were separately used to oxidize CD4 whereas H-XOOX was used to oxidize Xyl4. Cello- and xylo-tetramers served as reference cell wall oligomers with a proven elicitor activity in *A. thaliana* (Fernández-Calvo et al., 2024; Locci et al., 2019). The enzymatic reactions also occurred in the presence of a mammalian catalase, here used to recycle the hydrogen peroxide generated from oligosaccharide oxidation into H_2_O and O_2_, molecules both required for the oxidative reaction catalyzed by OSOXs (Pontiggia et al., 2020; Scortica et al., 2022). Following the enzymatic reactions, the oxidized tetra-saccharides were firstly checked by HPLC analysis and then were quantified by colorimetric assays (Fig. **1b,c**). In accordance with our previous analysis (Scortica et al., 2023), neutral oligosaccharides, once oxidized, strongly bound to the Pb-affinity column and, differently from their non-oxidized counterparts, could not be eluted by a isocratic gradient (Fig. **1b,c**-*left*). An ultra-filtration step allowed to separate OSOXs and catalase from the oxidized sugars that, in turn, were evaluated by two different colorimetric assays (Dubois et al., 1956; Lever, 1972). The quantification of total and reducing sugars performed on the three different oligomer preparations revealed an oxidation yield greater than 85-90% (Fig. **1b,c**-*right*). In conclusion, the pure OSOXs (Fig. **1a**) were successfully used to obtain two different mixtures of ox-CD4 [ox-CD4 (H-GOOX), ox-CD4 (FHS-CELLOX1)] and one mixture of ox-Xyl4 [ox-Xyl4 (H-XOOX)] (Fig. **1b,c**).

**Fig. 1.**
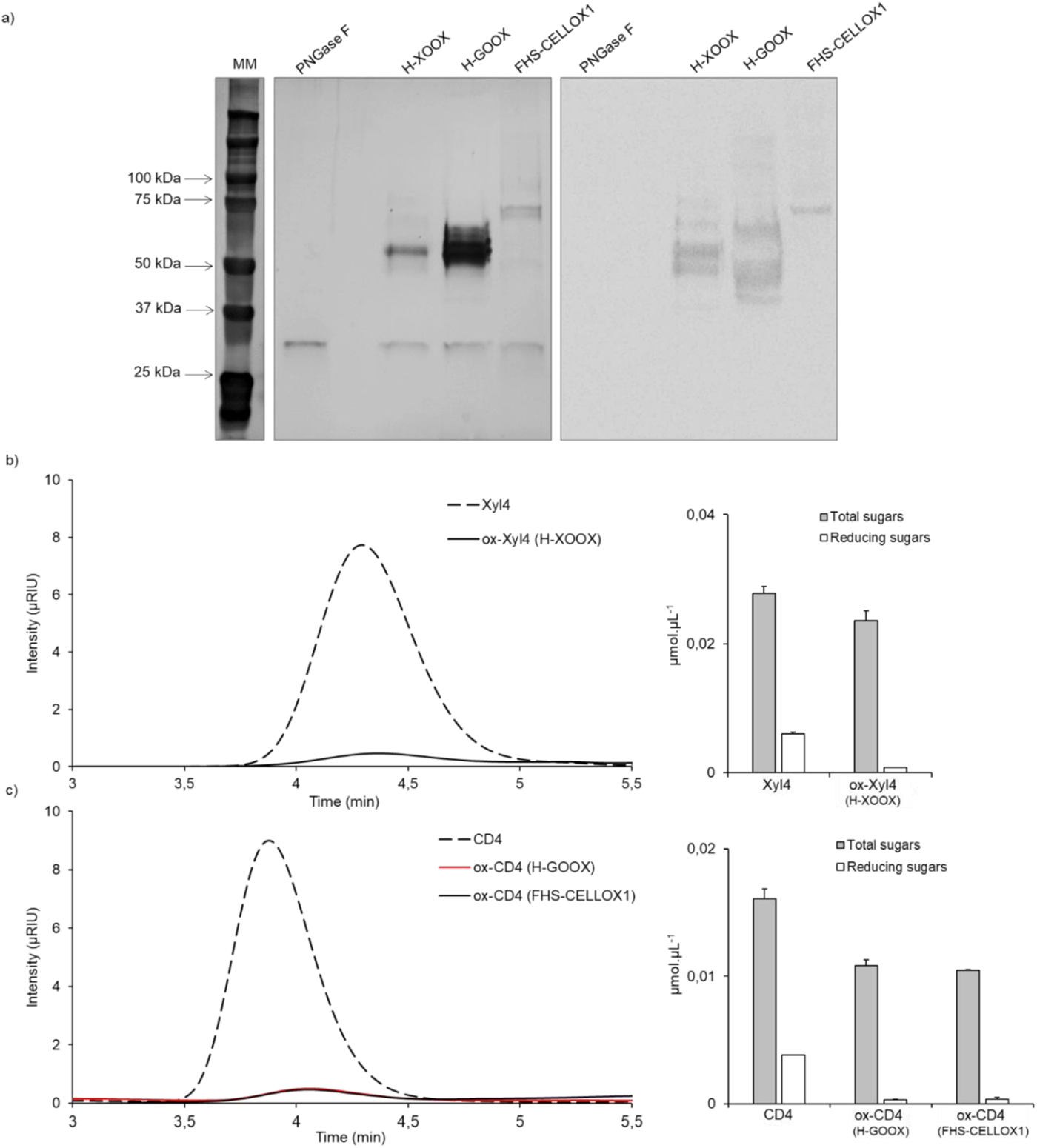
Oxidized oligosaccharides produced by recombinant plant and fungal OSOXs. a) *Left*, SDS-PAGE/silver nitrate staining analysis of purified OSOXs upon PNGase F treatment. Molecular weight marker (MM) is also reported. *Right*, immuno-decoration analysis of the same samples shown on the left by using an antibody directed against the 6xhis tag. B) *Left*, chromatographic analysis and (*right*) quantification of total and reducing sugars carried out on a Xyl4 preparation before (Xyl4) and after incubation with H-XOOX [ox-Xyl4 (H-XOOX)]. c) *Left*, chromatographic analysis and (*right*) quantification of total and reducing sugars carried out on a CD4 preparation before (CD4) and after incubation with H-GOOX [ox-CD4 (H-GOOX)] or with FHS-CELLOX1 [ox-CD4 (FHS-CELLOX1)]. All the enzymatic reactions were performed in 20 mM TRIS HCl pH 7.0 and 20 mM NaCl at room temperature. Xyl4 and CD4 (i.e., oligosaccharides without enzymes) were used as reference. Values are mean ± SD (N = 3). [CD4, cello-tetraose; FHS-CELLOX 1, flag-his-sumoylated cellodextrin oxidase 1; H-GOOX, his-gluco-oligosaccharide oxidase; H-XOOX, his-xylo-oligosaccharide oxidase; ox-CD4, oxidized cello-tetraose; ox-Xyl4, oxidized xylo-tetraose; PNGase F, Peptide:N-glycosidase F; Xyl4, xylo-tetraose].

### Plant defense responses are induced to a lesser extent by oxidized tetra-saccharides regardless of the type of OSOX used for their oxidation

To check whether ox-Xyl4 and ox-CD4 are still active as elicitors, we investigated three early defense responses (i.e., MAPK activation, extracellular accumulation of H_2_O_2_ and defense marker gene induction) on Arabidopsis seedlings treated with the oligosaccharide preparations shown in (Fig. **1b,c**). Here, the pathogen-associated molecular pattern (PAMP) “flg22” was used as positive control (Giovannoni, Lironi, et al., 2021; Giovannoni, Marti, et al., 2021) (Fig. **2**). The phosphorylation status of MPK3 and MPK6 in elicitor-treated seedlings was examined by an immuno-decoration analysis using the anti-pTpY antibody, which recognizes the double-phosphorylated activation loop of MAPKs in the TEY motif (Ichimura, 2002; Zhang & Dong, 2007). Compared to the treatment with Xyl4 and CD4, the phosphorylation status of MPK6 and MPK3 in seedlings treated with the oxidized tetra-saccharides was lower (ox-Xyl) or almost absent (ox-CD4), indicating that oxidized oligomers were worse elicitors than their non-oxidized counterparts (Fig. **2a**). Notably, the entity of such a decreased phosphorylation was similar in the seedlings treated with ox-CD4 (H-GOOX) and ox-CD4 (FHS-CELLOX1) (Fig. **2a**). The extracellular accumulation of H_2_O_2_ was measured in the culture medium of seedlings treated with the same oligosaccharide preparations by using the xylenol orange assay (Benedetti et al., 2018). Upon elicitor-treatment, the accumulation of H_2_O_2_ was 2-3-fold lower in the culture medium of seedlings treated with ox-Xyl4 and ox-CD4 (Fig. **2b**). Also in this case, the decrease of H_2_O_2_ accumulation was almost comparable in the seedlings treated with ox-CD4 (H-GOOX) and ox-CD4 (FHS-CELLOX1) (Fig. **2b**). Interestingly, this analysis also highlighted different kinetics of extracellular H_2_O_2_ accumulation between PAMP- and DAMP-treated plants since the extracellular accumulation of H_2_O_2_ was bi-phasic (30 and 90 min) and mono-phasic (30 min), respectively, for flg22- and oligosaccharide-treated seedlings (Fig. **2b**). The expression level of four defense-related genes known to be induced by OGs, CDs and flg22 was analyzed by qRT-PCR, namely *FOX1* (At1g26380), *CYP81F2* (AT5G57220), *FRK1* (AT2G19190) and *WRKY33* (AT2G38470) which generally peak at around 1 h after elicitation (Costantini et al., 2024; Galletti et al., 2008; Gravino et al., 2017) (Fig. **3**). In particular, *FOX1* encodes a BBE-l enzyme converting indole-cyanohydrin into the antimicrobial compound indole-3-carbonyl nitrile (Boudsocq et al., 2010; Rajniak et al., 2015), *CYP81F2* encodes a cytochrome P450 monooxygenase involved in the biosynthesis of indole glucosinolate (Asai et al., 2002; Nafisi et al., 2007), *FRK1* encodes a receptor-like protein kinase involved in early defense signaling whereas *WRKY33* encodes a transcription factor regulating the antagonistic relationship between the defense pathways induced by phytopathogens with different trophic styles. Compared to treatment with Xyl4 and CD4, plants treated with oxidized elicitors exhibited reduced induction of all investigated genes, confirming that oxidized oligosaccharides are less effective elicitors (Fig. **2-3**). This holds true regardless of their chemical composition (i.e., 1,4-β-linked-xylose or -glucose) or the OSOX used for their *in vitro* oxidation, whether fungal H-GOOX or plant FHS-CELLOX1 (Fig. **1**).

**Fig. 2.**
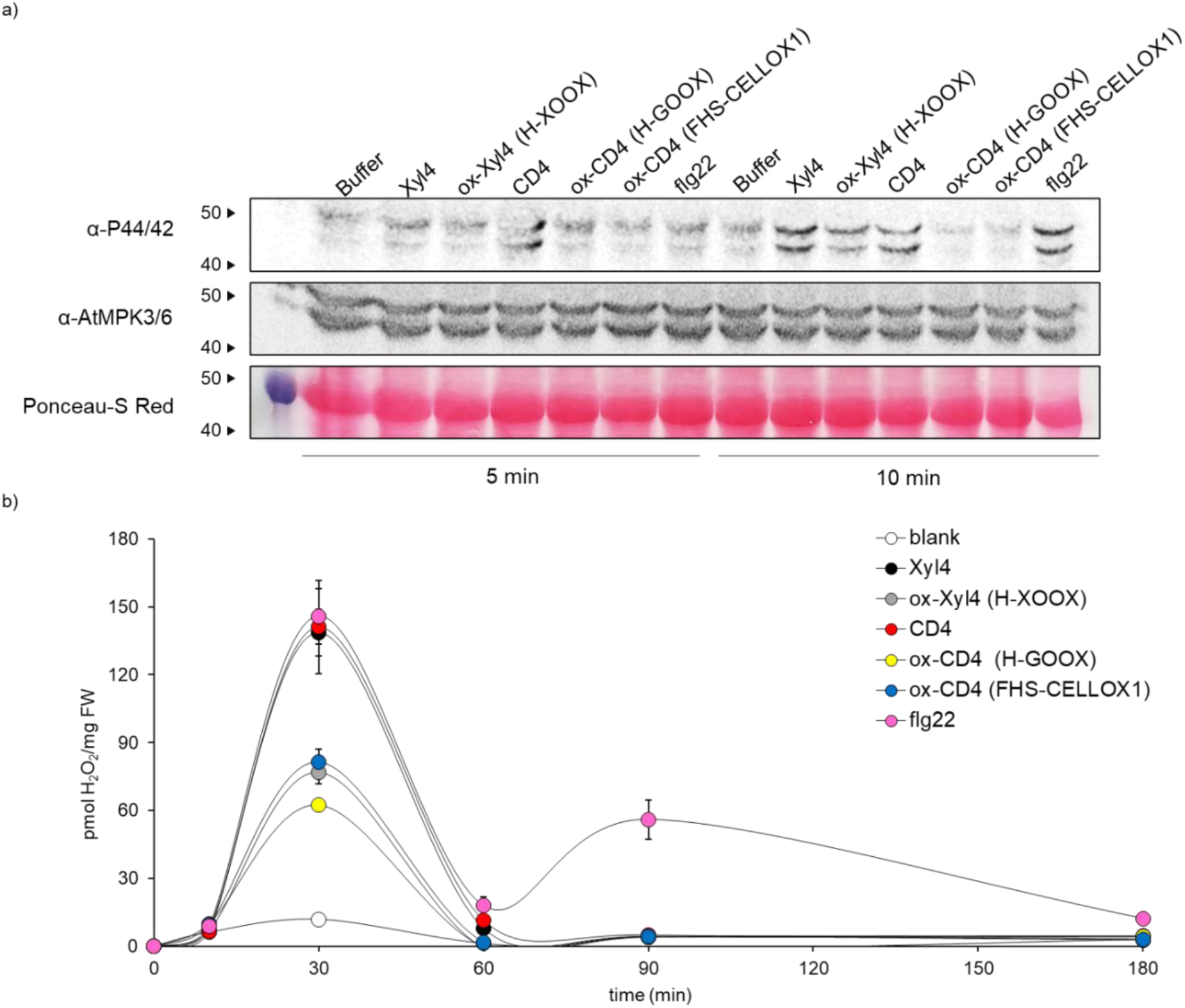
MAPK activation and extracellular H_2_O_2_ production are reduced in Arabidopsis seedlings treated with oxidized oligosaccharides. a) Immunoblot analysis on MPK3/6 phosphorylation in response to Xyl4, ox-Xyl4, CD4, ox-CD4 (H-GOOX), ox-CD4 (FHS-CELLOX1) and flg22. 10-day-old Col-0 seedlings were treated for 5 and 10 min with not oxidized and oxidized tetra-saccharides. Phosphorylated MPK3 and MPK6 (pMPK3/6) were detected by a α-phospho-p44/42 MAPK (Erk1/2) (Thr202/Tyr204) antibody. Total MPK3 and MPK6 were detected using α-AtMAPK3/6 antibodies. Ponceau-S Red staining was used to verify the equal loading. The experiment was repeated three times with similar results. b) H_2_O_2_ accumulation in the growth medium of wild-type seedlings after treatments with not oxidized and oxidized tetra-saccharides for the indicated time. Treatment with flg22 was used as positive control. Detection of H_2_O_2_ was performed by using the xylenol orange assay. Bars represent mean ± SD (n = 5). In all the experiments here reported, the treatment with the reaction buffer (blank) was used as negative control. [CD4, cello-tetraose; FHS-CELLOX 1, flag-his-sumoylated cellodextrin oxidase 1; fl22, flagellin 22 oligopeptide; H-GOOX, his-gluco-oligosaccharide oxidase; H-XOOX, his-xylo-oligosaccharide oxidase; ox-CD4, oxidized cello-tetraose; ox-Xyl4, oxidized xylo-tetraose; Xyl4, xylo-tetraose].

**Fig. 3.**
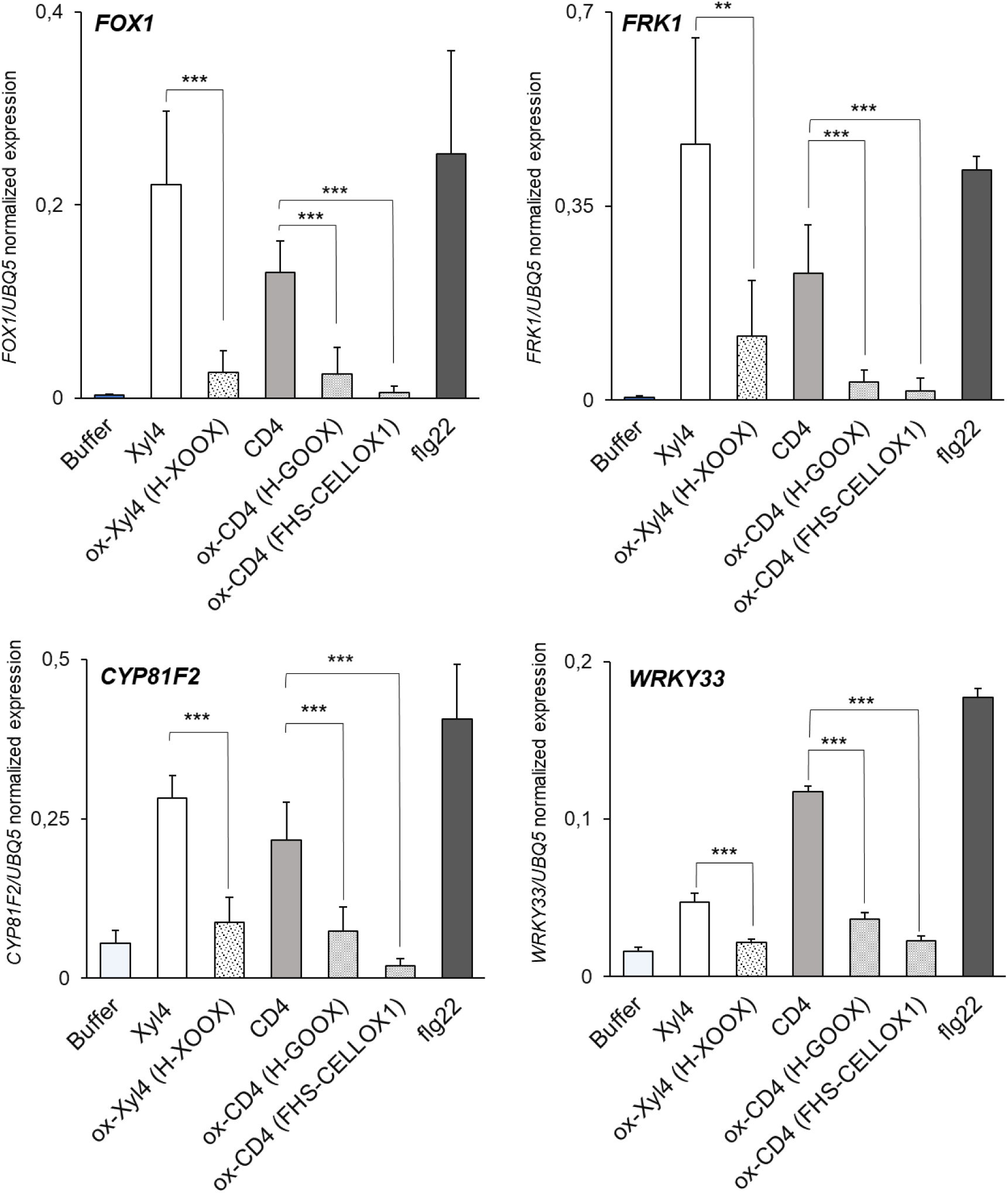
Induction of defense marker genes is reduced in Arabidopsis seedlings treated with oxidized oligosaccharides. Expression of the defense marker genes *FOX1*, *FRK1*, *CYP81F2* and *WRKY33* was evaluated by qRT-PCR in 10-day-old Col-0 seedlings after 1h-treatment with the only reaction buffer, Xyl4, ox-Xyl4 (H-XOOX), CD4, ox-CD4 (H-GOOX), ox-CD4 (FHS-CELLOX1) and flg22, the latter used as positive control. Normalization of gene expression was performed by using *UBQ5* a reference. Values are means of three independent biological experiments. Bars represent mean ± SD (n = 3). Asterisks indicate statistically significant differences between treatments with not oxidized oligosaccharides and their enzymatically oxidized counterparts, according to Student’s t-test (*, *p* < 0.05; **, *p* < 0.01; ***, *p* < 0 .001). [CD4, cello-tetraose; FHS-CELLOX 1, flag-his-sumoylated cellodextrin oxidase 1; fl22, flagellin 22 oligopeptide; H-GOOX, his-gluco-oligosaccharide oxidase; H-XOOX, his-xylo-oligosaccharide oxidase; ox-CD4, oxidized cello-tetraose; ox-Xyl4, oxidized xylo-tetraose; Xyl4, xylo-tetraose].

In conclusion, Xyl4 and CD4 triggered MAPK activation, extracellular H_2_O_2_ accumulation and defense marker gene induction in Arabidopsis seedlings, while ox-Xyl4 and ox-CD4 were weak or inactive as elicitors, regardless of the type of OSOX used for their *in vitro* oxidation. It is worth nothing that ox-CD4 and ox-Xyl4 retained some residual elicitor activity (Fig. **2-3**), which may be due to incomplete *in vitro* oxidation (Fig. **1b,c**-*right*).

### FHS-CELLOX1 and H-XOOX are characterized by a tetra-saccharide-dependent discoloration activity when acting on a [guaiacol: tetra-guaiacol] mixture

The existence of OSOXs in both plants and microbes makes the comprehension of their physiological role not easily inferable (Daniel et al., 2017; Pontiggia et al., 2020). *In vitro*, for example, the OSOX orthologs H-GOOX and FHS-CELLOX1 were both able to convert a DAMP (CD4) into a molecule with null elicitor activity (ox-CD4) in *A. thaliana* by concomitantly producing H_2_O_2_ (Fig. **1-3**). The scenario becomes even more complex when we consider that both plants and fungi express OSOXs during their interaction (Pontiggia et al., 2020). To further investigate this aspect, we analyzed the different redox propension of H-XOOX, H-GOOX and FHS-CELLOX1 in an attempt to highlight differences that were not evident in previous analyses (Costantini et al., 2024; Ferrari et al., 2016; Lee et al., 2005; Locci et al., 2019; Scortica et al., 2022; Scortica et al., 2023). Given that (i) the plant cell wall is primarily composed of polysaccharides and polyphenols, and (ii) OSOXs function in the extracellular environment, their ability to transfer electrons from cell wall oligosaccharides to plant phenolics was investigated. As potential phenolic electron acceptor substrate, we used tetra-guaiacol, a tetra-phenolic compound obtained by the POD-catalyzed oxidative polymerization of the coniferyl alcohol analogue “guaiacol” (Fig. **S1**). First, we determined the amount of each OSOX generating a similar H_2_O_2_ rate in the presence of equimolar concentration (50 µM) of appropriate tetra-saccharide (Fig. **4a**). Then, the same enzyme amounts were separately incubated in a reaction buffer containing a [guaiacol: tetra-guaiacol] mixture and POD. The assay, referred to as TG-reduction assay, included a concentration of tetra-guaiacol (∼150 µM) similar to that of dissolved oxygen in ultrapure water (180-380 µM). This was done to provide OSOXs with an equal likelihood of transferring electrons from the C1-end of each tetra-saccharide to either molecular O₂ or tetra-guaiacol (Gray, 2001). The simultaneous presence of guaiacol and tetra-guaiacol in the reaction, at an approximate molar ratio of 4:1, facilitated the identification of the redox propensity of each OSOX. This approach prevented the masking effect that could occur if only one phenolic species was present (Scortica et al., 2023). Surprisingly, the brownish color of the solution, typically attributed to tetra-guaiacol, decreased over time in the presence of two out of three OSOX/tetra-saccharide combinations (Fig. **S2**). In particular, the H-XOOX/Xyl4 and FHS-CELLOX/CD4 combinations displayed, respectively, a 8.6-fold and 1.3-fold higher discoloration activity than H_2_O_2_-generating activity, whereas the H-GOOX/CD4 combination did not display any discoloration effect (Fig. **4a,b**). Based on these results, we demonstrated that OSOXs with seemingly similar substrate specificities (i.e., H-GOOX and FHS-CELLOX1) actually catalyze different reactions.

**Fig. 4.**
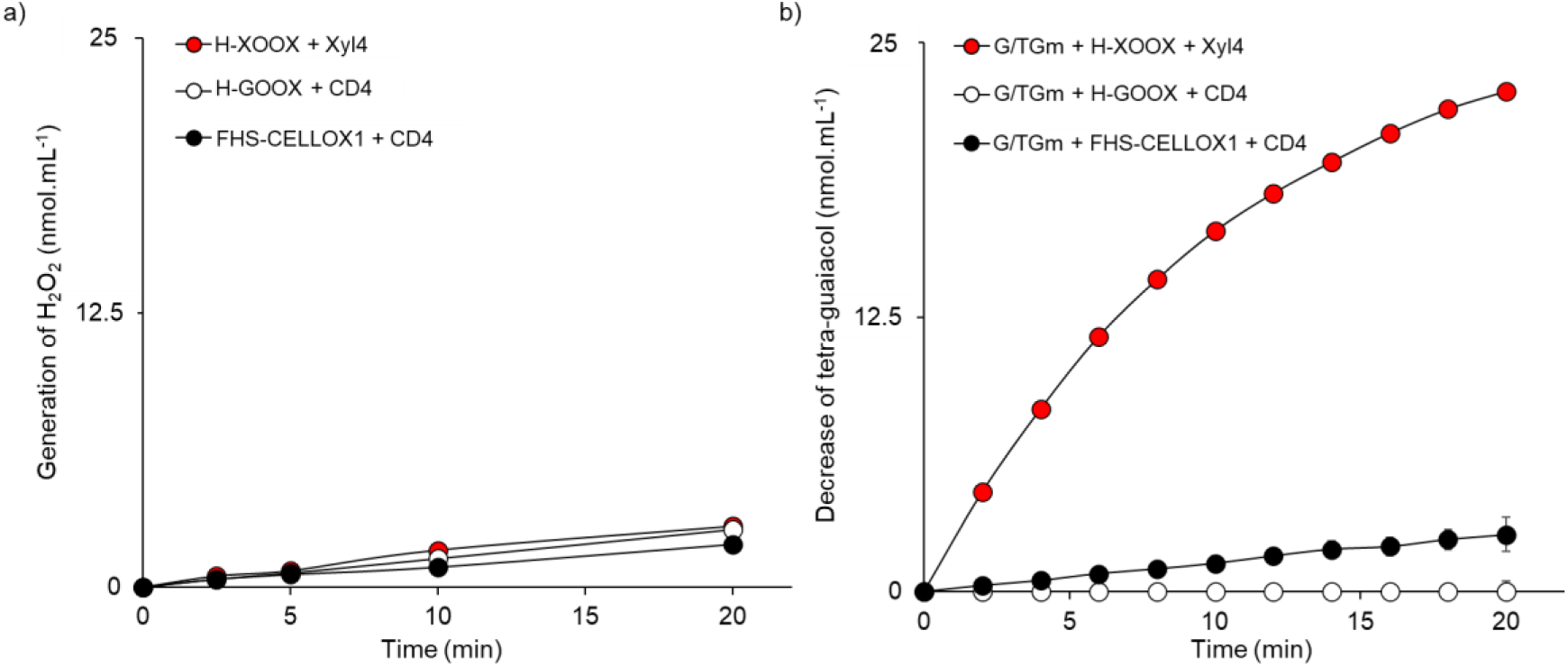
H_2_O_2_-generating and tetra-guaiacol-discoloration activity of different OSOX/oligosaccharide combinations. a) H_2_O_2_-generating and (b) tetra-guaiacol-discoloration activity, the latter expressed as decrease of tetra-guaiacol over time, at pH 5.0 by H-XOOX/Xyl4, H-GOOX/CD4 and FHS-CELLOX1/CD4 combinations as determined by the xylenol orange and TG-reduction assay, respectively. (a, b) The reactions were performed using different concentrations of H-XOOX (740 nM), H-GOOX (1.5 nM) and FHS-CELLOX1 (4 nM) in the presence of each appropriate tetra-saccharide (50 µM), and (b) a guaiacol/tetra-guaiacol mixture (G/TGm). Values are mean ± SD (N = 3). [CD4, cello-tetraose; FHS-CELLOX 1, flag-his-sumoylated cellodextrin oxidase 1; G/TGm, guaiacol/tetra-guaiacol mixture, H-GOOX, his-gluco-oligosaccharide oxidase; H-XOOX, his-xylo-oligosaccharide oxidase; OSOXs, oligosaccharide oxidases; Xyl4, xylo-tetraose].

### The tetra-saccharide dependent discoloration activity of OSOXs involved a scavenging effect against di- and tri-guaiacol radicals

The discoloration activity of H-XOOX/Xyl4 and FHS-CELLOX1/CD4 combinations was more pronounced in a reaction buffer predominantly composed of tetra-guaiacol, i.e. with a [guaiacol: tetra-guaiacol] ratio in favor of tetra-guaiacol (Movie S1). The absorption spectra obtained during these enzymatic reactions were consistent with our previous results (Fig. **4b**) but also provided additional insight (Fig. **5**). In the presence of H-XOOX/Xyl4 and FHS-CELLOX1/CD4 combinations, the characteristic double absorption peak of tetra-guaiacol (at 423 and 475 nm) was markedly reduced after 10 minutes of reaction. Concurrently, a new double absorption peak appeared at shorter wavelengths (266 and 292 nm) (Fig. **5**). This double absorption peak indicated that the reaction product(s) resulting from the discoloration activity of the OSOX/oligosaccharide pair were distinct from the starting substrate, i.e., guaiacol (which has a single absorption peak at 280 nm) (Fig. **5**). However, the addition of fresh H_2_O_2_ to the exhausted reaction (i.e., when the tetra-saccharides had been fully oxidized by OSOXs) restored the double absorption peak typically associated with tetra-guaiacol. This clearly demonstrated that the product(s) generated by OSOXs served as substrates for POD, and vice versa (Fig. **5**).

**Fig. 5.**
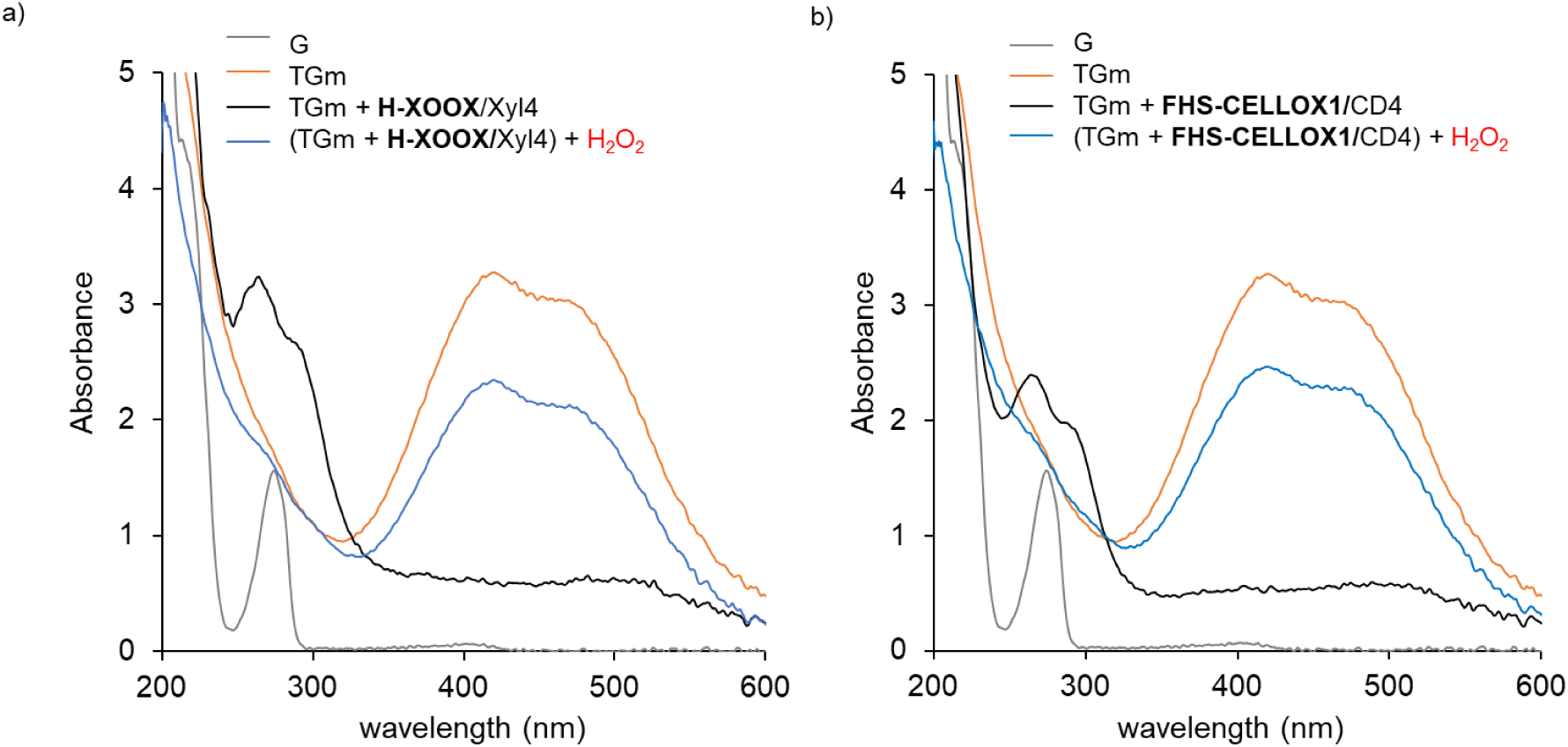
Absorption spectra of the tetra-guaiacol mixture during the reaction catalyzed by different OSOX/oligosaccharide combinations. Changes in the UV/Visible absorption of tetra-guaiacol mixture (TGm) after 10-minutes from the addition of a) H-XOOX/Xyl4 and b) FHS-CELLOX1/CD4, and after the addition of fresh H_2_O_2_ (600 µM) to the exhausted enzymatic reactions. TGm was obtained by adding H_2_O_2_ (600 µM) in the presence of guaiacol (600 µM) and POD (0.05 g.L^-1^), with the latter present in all the reactions here analyzed. The enzymatic reactions were performed in distilled water by adding the appropriate oligosaccharide (600 µM) and OSOX (20 ng.µL^-1^). The experiment was repeated three times with similar results. [CD4, cello-tetraose; FHS-CELLOX 1, flag-his-sumoylated cellodextrin oxidase 1; G, guaiacol; H-XOOX, his-xylo-oligosaccharide oxidase; OSOXs, oligosaccharide oxidases; POD, horseradish peroxidase type VI; TGm, tetra-guaiacol mixture; Xyl4, xylo-tetraose].

The same enzymatic reactions were also analyzed using a Q-TOF LC/MS system. Although this technique did not detect the guaiacol used as a reference, it successfully identified tetra-guaiacol as the [M + H]^+^ adduct with an m/z of 489 (Fig. **S3**). Surprisingly, di- and tri-guaiacol were also detected, but as diradical-like molecules, as evidenced by their reduced mass of -2 units less than their corresponding hydrogenated forms (i.e., m/z 245 *vs* 247 for di-guaiacol; m/z 367 *vs* 369 for tri-guaiacol) (Fig. **S3**). This suggested the presence of deprotonated hydroxylic group(s) in their structures (Samet et al., 2002). As expected, these molecular masses (m/z 245, 367 and 489) were specifically detected in the reaction containing guaiacol and POD only after the addition of H_2_O_2_ (Fig. **S3**). Furthermore, the extracted ion chromatograms (EICs) for the tri-guaiacol radical and tetra-guaiacol displayed distinct elution peaks, indicating the presence of different isomers for each oligomer (Fig. **S3b,c**-*upper panel*). Notably, the activity of XOOX/Xyl4 and FHS-CELLOX1/CD4 combinations on the mixture suppressed all these species in a size-dependent manner: di-guaiacol radical was suppressed more efficiently than tri-guaiacol radical, which was in turn suppressed more efficiently than tetra-guaiacol, with the latter being suppressed to a lesser extent (Fig. **6**). This trend may be attributed to the more favorable redox potential of smaller phenolic radicals, which might fit more effectively into the active site of OSOXs, thus allowing for more efficient suppression. These results corroborate recent findings reported by (Scortica et al., 2023), which demonstrated that different OSOX/short oligosaccharide combinations exhibit strong scavenging activity towards the radical cation ABTS^•+^. On the other hand, the missed detection of “suppressed/hydrogenated” di- and tri-guaiacol aligns with the Q-TOF LC/MS system’s inability to detect the guaiacol used as reference. At this stage, we could not clearly determine whether the suppression of di- and tri-guaiacol radicals by OSOXs involved only their scavenging or also their depolymerization into guaiacol (Fig. **S4**).

**Fig. 6.**
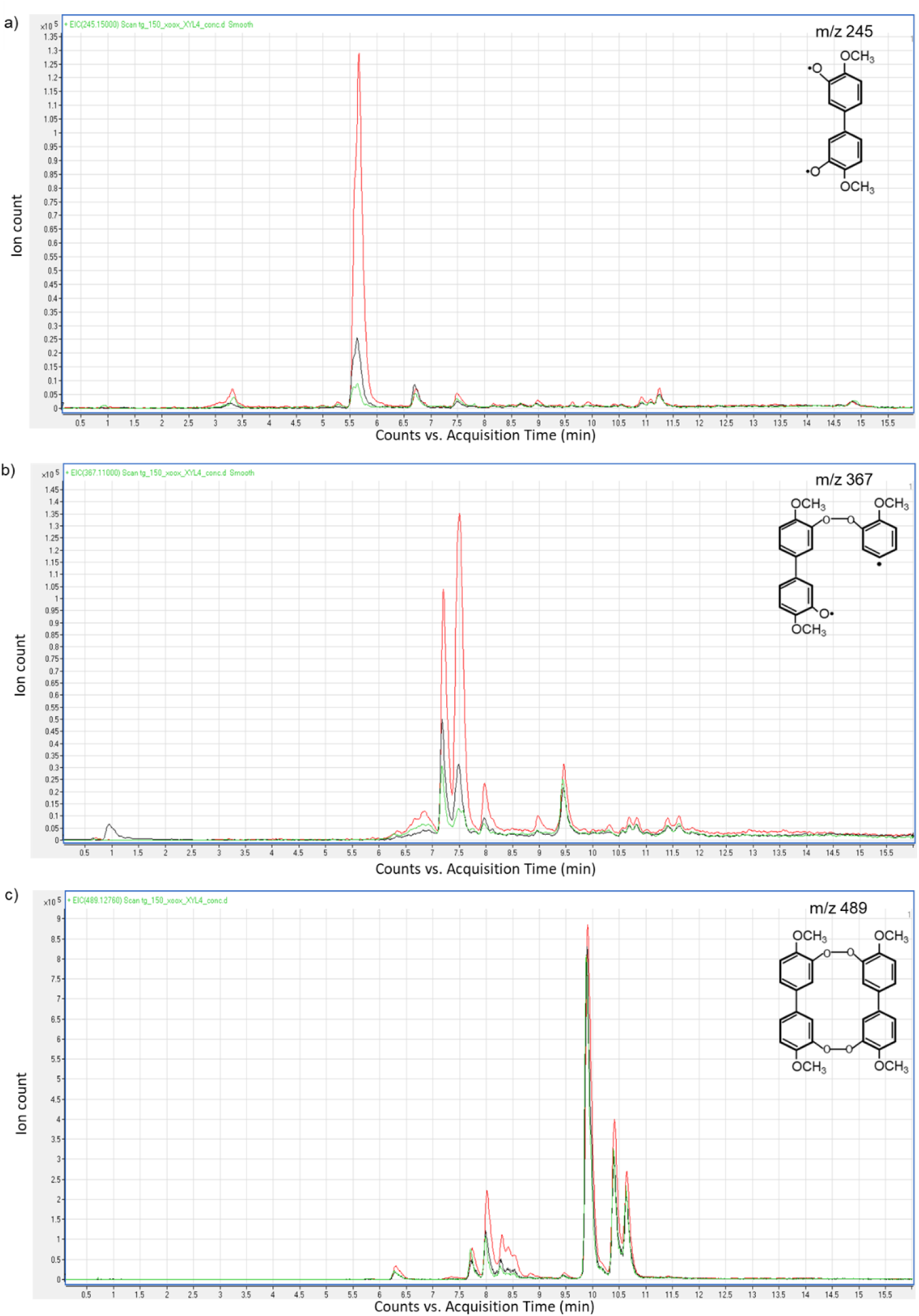
Decrease of three guaiacol-based oligomers upon the addition of two different OSOX/oligosaccharide combinations to the tetra-guaiacol mixture. EIC of three different ions as detected after 10 minutes from the addition of H-XOOX/Xyl4 (green line) and FHS-CELLOX1/CD4 (black line) combinations to the tetra-guaiacol mixture. The ion masses under investigation were characterized by a m/z of a) 245, b) 367 and c) 489. EIC of the same ions at the beginning of the reaction, i.e., without the addition of any OSOX/oligosaccharide combination, is reported as reference (red line). For each ion, a hypothetical structure is also reported. Mass ions were mainly detected as [H+M]^+^ adducts. [CD4, cello-tetraose; FHS-CELLOX 1, flag-his-sumoylated cellodextrin oxidase 1; H-XOOX, his-xylo-oligosaccharide oxidase; Xyl4, xylo-tetraose].

To deeper investigate this aspect, the same enzymatic reactions were also analyzed by HPLC chromatography. This technique was employed to measure the amount of guaiacol at three different time-points of enzymatic reaction, i.e., before and after the addition of H_2_O_2_, and 10 minutes after the addition of H-XOOX/Xyl4 and FHS-CELLOX1/CD4 combinations to the tetra-guaiacol mixture (Fig. **S5**). In the presence of H_2_O_2_, POD oxidized most of the starting guaiacol but the newly formed oligomers, such as tetra-guaiacol, were not clearly detected. The discoloration activity of H-XOOX/Xyl4 and FHS-CELLOX1/CD4 combinations on the tetra-guaiacol mixture slightly decreased the amount of residual guaiacol, an effect ascribable to a side-production of H_2_O_2_ by OSOXs and that POD, in turn, used to oxidize a small amount of the remaining guaiacol. This result excluded that OSOXs depolymerized di- and tri-guaiacol radicals into guaiacol (Fig. **S5**). However, the activity of H-XOOX/Xyl4 and FHS-CELLOX1/CD4 combinations also promoted a marked accumulation of an unknown compound(s) characterized by a double elution peak with a longer retention time (12.7-12.8 min) than guaiacol (8.6 min). After the addition of fresh H_2_O_2_ to the reaction, such compound(s) was quickly oxidized by POD (Fig. **S5**). This result agreed with all previous analyses and suggested that the compound(s) characterized by a double absorption peak (266 and 292 nm) shown in Figure 5 was the same unknown compound(s) of the HPLC analysis (Fig. **S5**), i.e., plausibly “suppressed/hydrogenated” di- and tri-guaiacol (Fig. **S4**).

By integrating the results obtained from the different analyses (UV/Visible absorption, Q-TOF LC/MS, HPLC), we concluded the discoloration activity of OSOXs mainly consisted of a scavenging effect against di- and tri-guaiacol radicals (Fig. **5-6**, Fig. **S4-5**, Movie **S1**). Moreover, we deduced that the brownish color of tetra-guaiacol mixture was mainly conferred by the presence of these radical species, precisely because the marked discoloration observed in the presence of different OSOX/oligosaccharide combinations was mainly associated to the suppression of di- and tri-guaiacol radicals than tetra-guaiacol (Movie **S1**, Fig. **6**).

## Discussion

We demonstrate that C1-oxidized cell wall oligosaccharides lose their ability to function as DAMPs in *A. thaliana*, and that their oxidation can be catalyzed by either microbial or plant OSOX. It is likely that the open-chain acid residue at the C1-end may impair binding with plant recognition receptors (PRRs), highlighting the reducing end as critical for oligosaccharide-receptor interactions. (Brutus et al., 2010; Fernández-Calvo et al., 2024). It is also possible that, in addition to PRR-mediated recognition, the signal transduction cascade triggered by a cell wall DAMP may require its oxidation. Notably, a mutant plant with a null response to a cell wall DAMP has never been isolated, supporting the idea that cell wall DAMP-mediated signaling involves multiple pathways. For example, DAMP perception through an OSOX-like redox reaction might allow plants to sense both the type (through the different substrate specificities of OSOXs) and amount of DAMP accumulated over time (through the release of a reactive product obtained from its oxidation, e.g., H_2_O_2_), enabling a defense response proportional to the entity of cell wall damage received (Scortica et al., 2022). It is worth noting that C1-oxidized oligosaccharides used to treat plants have been usually prepared *in vitro* (Benedetti et al., 2018; Costantini et al., 2024; Locci et al., 2019), preventing endogenous OSOXs from using them as an *in vivo* electron source. Consequently, plant response triggered by *in vitro*-oxidized oligosaccharides will inevitably be partial (Spiro et al., 1998; Benedetti et al., 2018; Locci et al., 2019; Costantini et al., 2024).

Although fungal H-GOOX and plant FHS-CELLOX1 have similar substrate specificity, meaning they can oxidize the same oligosaccharides, they exhibited different discoloration activities against a [guaiacol: tetra-guaiacol] mixture (Fig. **4a,b**); this result suggested that their physiological role is more related to the type of final electron acceptor substrate (e.g., O_2_ *vs* phenolic radicals) rather than to the common starting substrate acting as the electron donor (i.e., CD4).

In this context, transferring electrons from the reducing end of cell wall oligosaccharides to oligo-phenolic radicals can help prevent excessive accumulation of reactive oxygen species (ROS) in the extracellular environment. Notably, lignol radicals are formed not only by plant metallo-enzymes (e.g., peroxidases and laccases) for lignin synthesis and remodeling, but also by microbial enzymes to decompose lignin during pathogen attacks (Janusz et al., 2017). In the apoplast, both the synthesis and degradation of lignin involve the formation of similar reaction intermediates, i.e., phenolic radicals. This explains why both plant and microbial OSOXs are active on guaiacyl radical-based compounds. In a hypothetical multi-enzyme system, a CELLOX1/CD4 combination could suppress phenolic radicals in the apoplast, which can then be re-oxidized by a POD using extracellular hydrogen peroxide generated by other enzymatic systems (Fig. **5**). In line with this view, CELLOX1 may indirectly reduce the total amount of ROS produced during infection by recycling substrates that can be oxidized by PODs, thereby consuming H_2_O_2_ (Fig. **5**, Fig. **S5**) (Costantini et al., 2024; Locci et al., 2019; Scortica et al., 2023). Alternatively, xylo-oligosaccharides produced by xylanolytic fungi could serve as an electron source for XOOX to counteract lignin polymerization catalyzed by plant PODs.

In our experimental conditions, fungal GOOX, unlike the plant ortholog CELLOX1, was inactive against the guaiacol/tetra-guaiacol mixture. Probably, H_2_O_2_-generating and discoloration activities of GOOX were almost comparable in the guaiacol/tetra-guaiacol mixture, resulting in no clear net activity (Fig. **4b**, Fig. **S2b**). To understand the biological significance of this result, it is worth noting that the virulence of certain fungal pathogens, such as *Botrytis cinerea*, is enhanced by the accumulation of ROS in plant tissues. Indeed, these pathogens possess various H_2_O_2_-generating systems to sustain the infection process (Rolke et al., 2004). In this scenario, a fungal OSOX with higher scavenging activity than H_2_O_2_-generating activity may be counter-productive for a successful infection, indicating that the function of each fungal OSOX is closely related to the specific lifestyle and infection strategy of its microbial producer.

Fungal oligosaccharide oxidases belong to the CAZy (Carbohydrate Active Enzymes database, http://www.cazy.org/) AA7 family (Auxiliary Activity Family 7) and feature a bi-covalently attached FAD cofactor (Drula et al., 2022). Similarly, plant BBE-l enzymes also share these structural characteristics, although they are currently not assigned into AA7 family (Momeni et al., 2021). It is noteworthy however, that the type of active site and oxygen reactivity motif are not identical between XOOX, GOOX and CELLOX1 (Fig. **S6**). A deeper biochemical and structural characterization of plant OSOXs will be fundamental to identify natural compounds acting as electron acceptors *in vivo*. In enzymology, oxidation by a FAD-dependent enzyme follows a Ping-Pong Bi-Bi mechanism: one substrate enters the active site, transfers a hydride to the flavin, and then is released. Subsequently, a second molecule forms a complex to re-oxidize the flavin (Daniel et al., 2017). However, the radical cation scavenging activity of OGOX1 occurs only in the presence of short OGs, indicating that the substrate steric hindrance, along with their redox potential, influences the dual scavenging/oxidase activity of OSOXs (Scortica et al., 2023).

## Conclusions

The trade-off between growth and defense involves allocating metabolic energy to maintain optimal health and fitness. Currently, the molecular mechanisms that enable plants to modulate defense and growth processes in response to cell wall damage are not fully understood. In this work, a novel role for OSOXs as metabolic connectors between glucidic and phenolic components of plant cell wall has been uncovered. OSOXs with similar substrate specificities can catalyze effectively diverse reactions due to their different abilities in redirecting the reducing power from cell wall oligosaccharides to phenolic radicals. The latter is an additional sub-functionalization feature that might explain the co-presence of vary OSOX paralogs in the same organism (Daniel et al., 2017; Pontiggia et al., 2020). Our analysis revealed that OSOXs, acting downstream of glycoside hydrolases, can either enhance or counteract the activity of metallo-enzymes on phenolic radicals. This finding suggests the existence of a complex network of enzymatic activities that has not been fully clarified so far. OSOXs are central players in the mechanisms evolved by plants to manage the oxidative stress caused by DAMP-triggered immunity. Advances in understanding these processes could lead to novel strategies for optimizing plant biomass exploitation in the biofuel industry and for mitigating disease symptoms associated with saccharide-based DAMPs and free radicals in animal tissues.

## CRediT authorship contribution statement

**Moira Giovannoni**: writing – original draft, visualization, methodology, investigation, formal analysis, data curation. **Anna Scortica**: writing – original draft, visualization, methodology, investigation, formal analysis, data curation. **Valentina Scafati**: visualization, methodology, investigation. **Emilia Piccirilli**: visualization, methodology, investigation. **Daniela Sorio**: visualization, methodology, formal analysis. **Manuel Benedetti**: writing – review & editing, writing – original draft, visualization, supervision, methodology, investigation, formal analysis, data curation, conceptualization, funding acquisition, project management. **Benedetta Mattei**: writing – review & editing, supervision, methodology, formal analysis, funding acquisition, project management.

## Acknowledgements

The authors gratefully acknowledge “Centro Piattaforme Tecnologiche” (University of Verona, Italy) for support and assistance in mass spectrometry analysis. This work was supported by the Italian Ministry of University and Research (PRIN2022 – PNRR P2022TPZW3, funded to M.B.; PRIN 2022WLZ4HB and PRIN2022 – PNRR P2022KLXEC, both funded to B.M.) and the University of L’Aquila (Progetti di Ateneo 2023 – H_2_O_2_-BBE, funded to M.B.).

