## Supplementary Material for "The reducing end of cell wall oligosaccharides is critical for DAMP activity in *Arabidopsis thaliana* and can be exploited by oligosaccharide oxidases in the scavenging of phenolic radicals"

### List of content

**Fig. S1.** Reaction scheme of the oxidative polymerization of guaiacol as catalyzed by POD.

**Fig. S2.** Measurement of discoloration activity by using different combinations of OSOXs and substrates.

**Fig. S3.** Ions specifically detected in the starting tetra-guaiacol mixture.

**Fig. S4.** Possible reactions involved in the discoloration activity of H-XOOX/Xyl4 combination on a guaiacol-based radical intermediate.

**Fig. S5.** Determination of guaiacol in the tetra-guaiacol mixture after the addition of different OSOX/oligosaccharide combinations.

**Fig. S6.** Multiple amino acid alignment between H-XOOX, H-GOOX and FHS-CELLOX1.

**Method S1.** Heterologous expression of plant and fungal OSOXs in *Pichia pastoris.*

**Method S2.** Plant materials and elicitors.

**Method S3.** Plant growth conditions.

**Method S4.** Determination of extracellular $H_2O_2$ accumulation.

**Method S5.** Evaluation of OSOX activity by xylenol orange and tetra-guaiacol reduction assays.

**Method S6.** Analysis of substrates/products in the enzymatic reactions by UV/Visible absorption, and Q-TOF LC/MS and HPLC systems.

**Method S7.** Bioinformatic analyses.

**Table S1.** Primers used in this work.

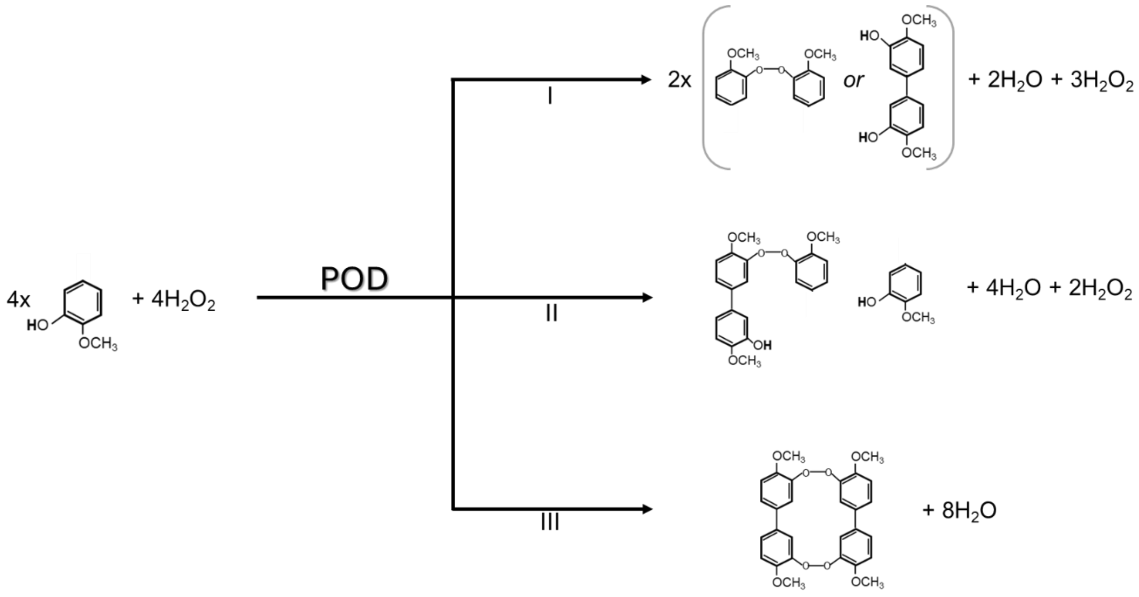


**Fig. S1. Reaction scheme of the oxidative polymerization of guaiacol as catalyzed by POD**. In the reaction scheme, the partial (I, II) and full (III) oxidative polymerization of guaiacol into the respective reaction intermediates (di-guaiacol and tri-guaiacol) and end-product (tetra-guaiacol) is shown. [POD: peroxidase].

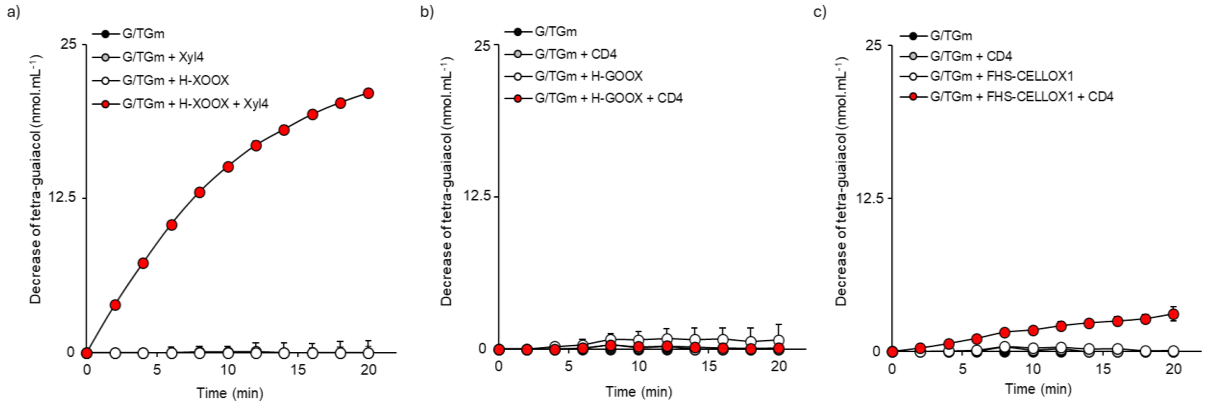


**Fig. S2. Measurement of discoloration activity by using different combinations of OSOXs and substrates.** Discoloration activity, expressed as decrease of tetra-guaiacol over time, at pH 5.0 by the activity of a) H-XOOX, b) H-GOOX and c) FHS-CELLOX1 as determined by the TG-reduction assay. Different amounts of OSOXs generating a comparable $H_2O_2$ rate were used (*see* Fig. **4a**) in the presence of different combinations of substrates. Values are mean ± SD (N=3). [CD4, cello-tetraose; FHS-CELLOX 1, flag-his-sumoylated cellodextrin oxidase 1; G/TGm, guaiacol/tetra-guaiacol mixture; H-GOOX, his-gluco-oligosaccharide oxidase; H-XOOX, his-xylo-oligosaccharide oxidase; Xyl4, xylo-tetraose].

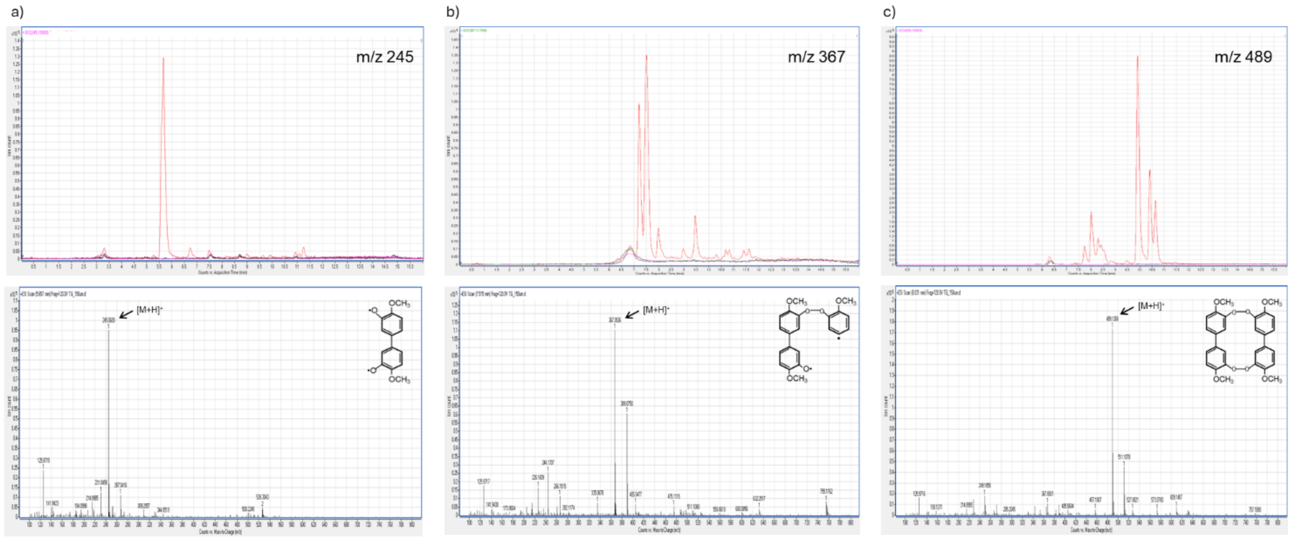


**Fig. S3. Ions specifically detected in the starting tetra-guaiacol mixture.** *Upper panel,* extracted ion chromatogram (EIC) of three different ions as detected upon addition of $H_2O_2$ to guaiacol (black line), to POD (pink line), and to [guaiacol + POD] (red line), the latter referred to as tetra-guaiacol mixture. EIC of the same ions detected in the sample [guaiacol + POD] without the addition of $H_2O_2$ (green line) is also reported. *Lower panel*, mass spectra of ions specifically detected in the tetra-guaiacol mixture [i.e., $H_2O_2$ + (guaiacol + POD)] were characterized by a m/z of a) 245, b) 367 and c) 489. For each ion, a hypothetical structure is also reported. Mass ions were mainly detected as $[M+H]^+$ adducts. [POD, horseradish peroxidase type VI].

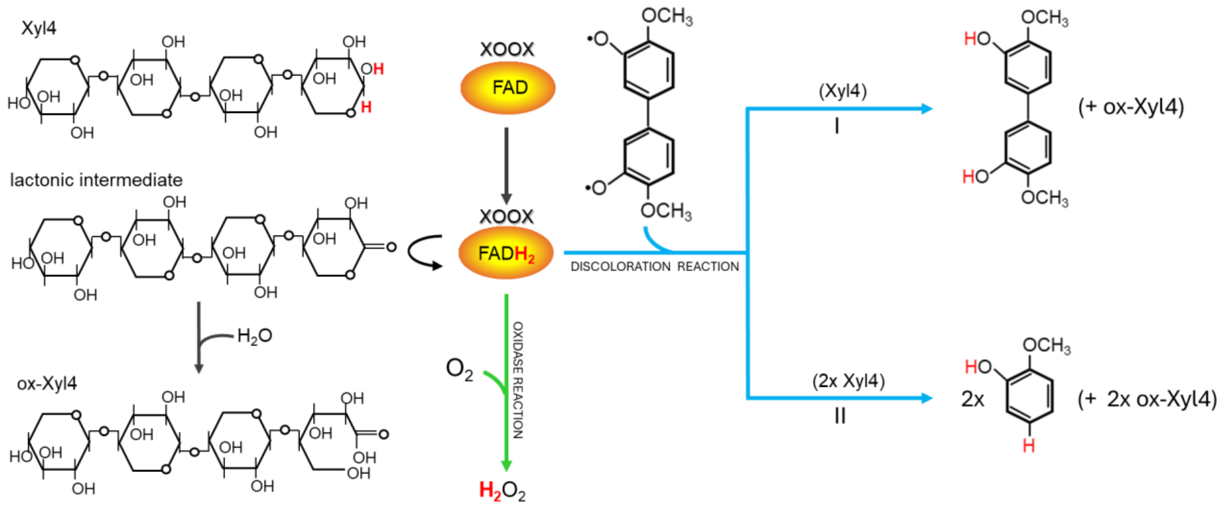


**Fig. S4. Possible reactions involved in the discoloration activity of H-XOOX/Xyl4 combination on a guaiacol-based radical intermediate.** Model of the dual oxidase/discoloration activity of H-XOOX/Xyl4 combination, here used as representative OSOX/oligosaccharide combination. In the oxidase reaction (green arrow), XOOX oxidizes Xyl4 to a lactonic intermediate by transferring two hydrides to molecular $O_2$ that, in turn, is converted into $H_2O_2$. In the discoloration reaction (turquoise arrow), XOOX transfers the two hydrides, instead of to molecular $O_2$, to di-guaiacol radical. The activity of H-XOOX/Xyl4 combination on a di-guaiacol radical can result in a reaction of scavenging (I) and/or depolymerization (II). In all reactions, the lactonic intermediate is then spontaneously hydrolyzed to xylo-tetraonic acid (ox-Xyl4). H-XOOX/Xyl4 and di-guaiacol radical are reported as representative OSOX/oligosaccharide combination and guaiacol-radical intermediate, respectively. The hydrogens involved in the reaction are reported in red. [H-XOOX, his-xylo-oligosaccharide oxidase; ox-Xyl4, oxidized xylo-tetraose; Xyl4, xylo-tetraose].

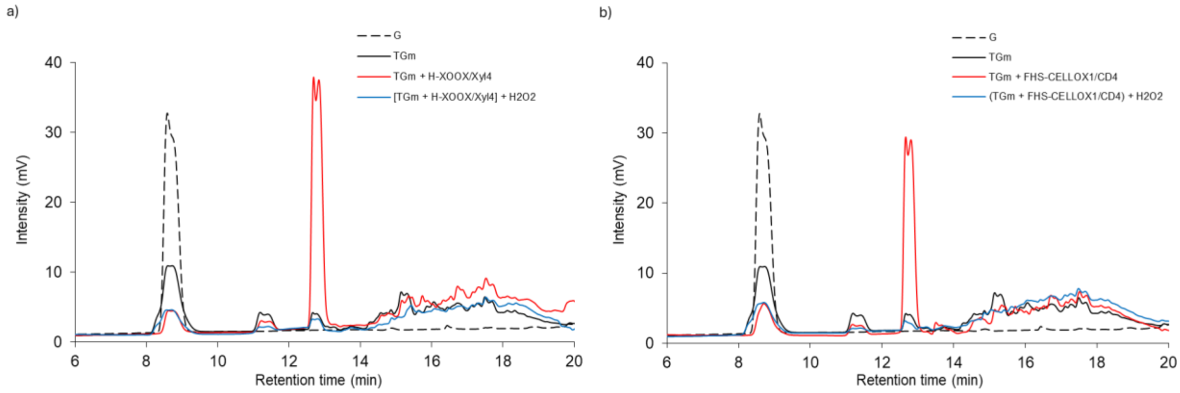


**Fig. S5. Determination of guaiacol in the tetra-guaiacol mixture after the addition of different OSOX/oligosaccharide combinations.** HPLC analysis on the reaction mixture: before (G) and after the addition of $H_2O_2$ (TGm), 10 minutes after the addition of a) H-XOOX/Xyl4 and b) FHS-CELLOX1/CD4 combinations to TGm, and after the addition of fresh $H_2O_2$ (600 µM) to the respective exhausted enzymatic reactions. TGm was obtained by adding $H_2O_2$ (600 µM) in the presence of guaiacol (600 µM) and POD (0.05 $g.L^{-1}$), with the latter present in all the reactions here analyzed. The enzymatic reactions were performed in distilled water by adding the appropriate oligosaccharide (600 µM) and OSOX (20 $ng.µL^{-1}$). The experiment was repeated three times with similar results. [CD4, cello-tetraose; FHS-CELLOX 1, flag-his-sumoylated cellodextrin oxidase 1; G, guaiacol; H-XOOX, his-xylo-oligosaccharide oxidase; POD, horseradish peroxidase type VI; TGm, tetra-guaiacol mixture; Xyl4, xylo-tetraose].

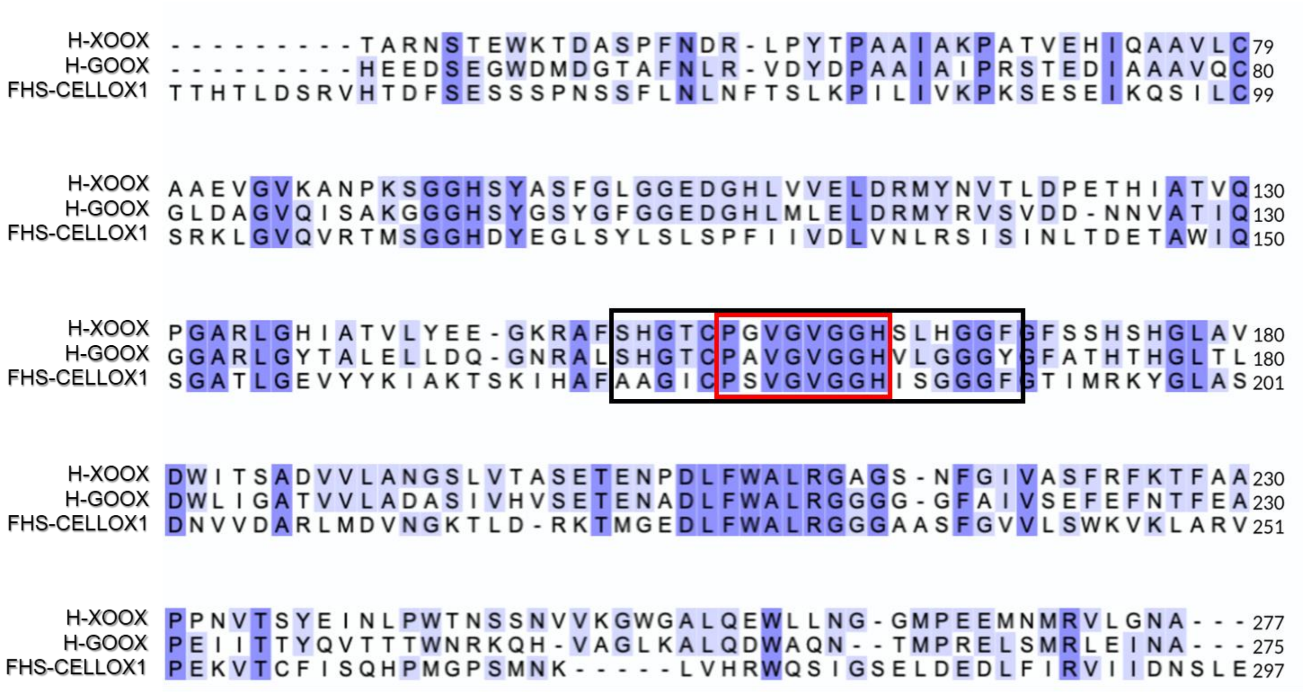


**Fig. S6. Multiple amino acid alignment between H-XOOX, H-GOOX and FHS-CELLOX1**. Intense violet indicates a residue conserved in all OSOXs under investigation, light violet indicates a residue conserved in two out of three OSOXs whereas a white background indicates absence of conserved residues. A black box indicates the "oxygen reactivity motif" whereas a red box indicates the "oxygen binding pocket" [P(G/A/S)VGVGGH]. Numbering is from the first amino acid of each mature protein. [FHS-CELLOX 1, flag-his-sumoylated cellodextrin oxidase 1; H-GOOX, his-gluco-oligosaccharide oxidase; H-XOOX, his-xylo-oligosaccharide oxidase].

**Method S1. Heterologous expression of plant and fungal OSOXs in *Pichia pastoris.***

The constructs pPICZαB/FHS-CELLOX1, pPICZαB/H-XOOX and pPICZαB/H-GOOX were transformed in *Escherichia coli* DH5α for plasmid propagation. After amplification, the constructs were linearized by SacI and transformed in *P. pastoris* by electroporation according to (Wu & Letchworth, 2004). Transformants were selected on solid YPDS medium [1% (w/v) yeast extract, 2% (w/v) peptone, 2% (w/v) dextrose, 1 M sorbitol] using zeocin as selection marker (100 $\mu g.mL^{-1}$). For protein expression at small scale, colonies of *P. pastoris* transformants were inoculated in 5 mL of YPD medium supplemented with zeocin (100 $\mu g.mL^{-1}$) and incubated at 28 °C in a rotary shaker at 180 rpm for 72h. After reaching the stationary growth phase, the culture was centrifuged and the pellet resuspended in 1.5 mL of Buffered Minimal Medium [BMM; 0.1 M K-phosphate (pH 6.0), 1.34% (w/v) YNB, 4 × $10^{-5}$% (w/v) biotin and 0.5% (v/v) methanol] to induce the expression of the fungal OSOXs and the growth prolonged for additional 72 h. The filtrates from 72 h-grown cultures were centrifuged and evaluated by SDS-PAGE and immune-decoration analysis by using a monoclonal anti-HIS antibody (AbHis, Bio-rad) to detect the expression of H-XOOX and H-GOOX (data not shown). In parallel, the oxidizing activities of H-XOOX and H-GOOX towards, respectively, Xyl4 and CD4 were analyzed in the same filtrates using the xylenol orange assay (Gay et al., 1999) (data not shown). At large scale, the purification of H-XOOX was achieved from a 72h-grown BMM culture. After cell removal by centrifugation (2500×g, 5 min), the culture filtrate was filtered with a sterile Polyether Sulfone (PES) filter (cut-off 0.2 μm), and then concentrated and dialyzed against a buffer composed of (50 mM Tris-HCl pH 7.5, 500 mM NaCl, 1 mM 2-mercaptoethanol and 10 mM imidazole) by using a modular tangential flow system (Vivaflow® 200, cut-off: 30 kDa). The dialyzed filtrate was loaded onto a HisTrap HP column (ThermoFisher, Waltham, USA) and the bound proteins were eluted at high imidazole concentration (500 mM). The eluted fractions containing H-XOOX were pooled and dialyzed in 50 mM Tris–HCl pH 7.0 and 100 mM NaCl. The dialyzed H-XOOX preparation was quantified by UV–visible spectrum of FAD ($\varepsilon 450nm = 11.300\ mM^{-1}.cm^{-1}$) and then analyzed by SDSPAGE/Silver nitrate staining. As already described by (Lee et al., 2005), H-GOOX failed to bind to the Ni-affinity column. Therefore, H-GOOX purification was performed by using hydrophobic interaction chromatography (HIC) carried out at pH 7.5 using HiTrap Phenyl FF-High Sub columns (GE Healthcare). Before HIC, $(NH_4)_2SO_4$ was added to the filtrate up to reach a final concentration of 2 M, then the sample was loaded onto a HiTrap Phenyl FF column pre-equilibrated with 50 mM Tris-HCl (pH 7.5) and 2M $(NH_4)_2SO_4$. The elution was performed by a linear gradient of a buffer constituted of 50 mM Tris-HCl (pH 7.5). Fractions containing the highest amount of H-GOOX were pooled, concentrated and then dialyzed in 50 mM Tris-HCl pH 7.5 and 50 mM NaCl. The amount of the purified H-GOOX was measured by using UV/Visible spectrum of FAD ($\varepsilon450nm = 11.300\ mM^{-1}.cm^{-1}$). The purified sample was also analyzed by SDS PAGE/ Silver nitrate staining. The activity of pure H-XOOX and H-GOOX was evaluated by using the xylenol orange assay using Xyl4 and CD4 as respective substrates. Transformation, selection of Pichia transformants and purification of FHS-CELLOX1 were performed by following the same procedures described in (Scortica et al., 2022).

**Method S2. Plant materials and elicitors.**

The *Arabidopsis thaliana* ecotype Columbia-0 (Col-0) seeds were used as wild-type plant in this study. The elicitors used in this study are xylo-tetraose (Xyl4, Megazyme, cod. O-XTE), cello-tetraose (CD4, Megazyme, cod. O-CTE) and flagellin 22 oligopeptide (flg22, EZBiolab, cod. 7201).

**Method S3. Plant growth conditions.**

The Fitotron® SGC2 was used as plant growth chamber (Weiss Technik). Plants grown at 22°C and 60% relative humidity with 16/8 h light/dark photoperiod and a light intensity of 120 $\mu mol.m^{-2}.sec^{-1}$. For seedling assays, seeds were surface sterilized using a sterilizing solution [0.01% w/v sodium dodecyl sulphate (SDS) and 1.6% v/v sodium hypochlorite (NaClO)] for 10 minutes, before being washed with autoclaved water. Later, ten seeds per well were grown in 2 mL of 0.5X Murashige and Skoog (MS) medium, pH 5.7, supplemented with 0.5% sucrose in multi-well plate system.

**Method S4. Determination of extracellular $H_2O_2$ accumulation.**

The amount of $H_2O_2$ produced during incubation of wild type seedlings (Col-0) with elicitors was performed using the xylenol orange assay (Jiang et al., 1990; Scortica et al., 2023). The $H_2O_2$ accumulation in the culture medium was measured in ten-day-old seedlings treated with each elicitor (80 μg.mL$^{-1}$) and water. The absorbance at 560 nm was measured at 0 (no treatment), 10, 30, 60, 90, 180 min after incubation with elicitors. Data were represented as pmol $H_2O_2$.mg$^{-1}$ FW seedlings * mL.

**Method S5. Evaluation of OSOX activity by xylenol orange and tetra-guaiacol reduction assays.**

$H_2O_2$-producing activity of different OSOX/oligosaccharide combinations was measured by the xylenol orange assay (Gay et al., 1999). Production of $H_2O_2$ was evaluated in 50 mM Na-Acetate pH 5.0 and 50 mM NaCl (0.1 mL) by adding each appropriate oligosaccharide (50 μM) in the presence of different amounts of H-XOOX, H-GOOX and FHS-CELLOX1. To produce a similar $H_2O_2$ rate under these reaction conditions, the pure OSOXs were used at 740 nM (H-XOOX), 1.5 nM (H-GOOX) and 4 nM (FHS-CELLOX1). Then, the xylenol orange assay was performed as previously reported (Scortica et al., 2023). The discoloration activity of H-XOOX/Xyl4, H-GOOX/CD4 and FHS-CELLOX1/CD4 combinations against a [guaiacol: tetra-guaiacol] mixture (G/TGm) was evaluated by using the tetra-guaiacol-reduction assay, hereafter referred to as TG-reduction assay. The basal mixture was prepared by adding $H_2O_2$ (600 μM) to a solution (50 mM Na-acetate pH 5.0 and 50 mM NaCl, 0.2 mL) containing guaiacol (1.2 mM) (Sigma-Aldrich) and horseradish peroxidase type VI (POD, 0.05 g.L$^{-1}$) (Sigma-Aldrich). After few seconds from the addition of $H_2O_2$, the color of this solution changes from colorless to brownish, an effect commonly attributed to the oxidative polymerization of guaiacol into tetra-guaiacol as catalyzed by POD. Based on our calculations, the basal mixture of TG-reduction assay was characterized by an approximate [guaiacol: tetra-guaiacol] molar ratio of 4:1; then, the precise amount of tetra-guaiacol was determined by measuring the absorbance at 470 nm ($\varepsilon$470nm = 26.6 mM$^{-1}$.cm$^{-1}$) in accordance with conventional accepted procedures. After the preparation of tetra-guaiacol, the enzymatic reactions (0.2 mL) were started by adding the different OSOX/oligosaccharide combinations. The assay was performed under the same conditions of temperature, tetra-saccharide and enzyme concentrations as those used in the xylenol orange assay. Reduction of tetra-guaiacol was measured by following the decrease of absorbance at 470 nm in continuous mode for 20 minutes. Values were subtracted to that of the control reaction (basal mixture + corresponding OSOXs) and then converted into nmol of tetra-guaiacol (ε470nm = 26.6 $mM^{-1}.cm^{-1}$). All the analyses were performed in triplicates by using an Infinite R M Nano200 spectrophotometer (Tecan AG, Mannedorf, Switzerland).

**Method S6. Analysis of substrates/products in the enzymatic reactions by UV/Visible absorption, and Q-TOF LC/MS and HPLC systems.**

For the acquisition of UV/Visible and Q-TOF LC/MS spectra, a modified version of TG-reduction assay was used. Here, the basal mixture, mainly composed of tetra-guaiacol (approximately 150 µM), was obtained by adding $H_2O_2$ (600 µM) to distilled water (0.2 mL) containing guaiacol (600 µM) and horseradish peroxidase type VI (POD, 0.05 $g.L^{-1}$). As observed for the TG-reduction assay, after few seconds from the addition of $H_2O_2$, the color of this solution also changes from colorless to brownish. Then, the reactions were completed by adding the appropriate oligosaccharide (600 µM) and OSOX (20 $ng.\mu L^{-1}$) to such mixture. The reactions containing [guaiacol + POD] and [tetra-guaiacol + POD, i.e., TGm] were used as control reactions. Changes in the UV/Visible absorption of tetra-guaiacol during TG-reduction assay were recorded by using NanoDrop™ One/One$^{C}$ (Thermofisher). Absorption spectra of enzymatic and control reactions were recorded after 10 minutes of reaction. For the Q-TOF LC/MS analysis, the same reactions were analyzed by using a 1260 HPLC system (Agilent Technologies, Waldbronn, Germany) in tandem with a Q-TOF mass spectrometer. For the separation of molecules, each sample (5 µL) was loaded onto a Zorbax Eclipse XDB (2.1 × 150 mm, 5 µm particle size, Agilent Technologies). The composition of mobile phase was as follows: 0.1% (v/v) formic acid (solvent A) and 0.1% (v/v) formic acid in acetonitrile (solvent B). The samples were eluted using a 19 min linear gradient at flow rate of 0.5 mL.min$^{-1}$ (hold at 20% for 2 min, from 20% to 100% solvent B in 13 min, hold at 100% for 2 min, hold at 20% for 4 min). The MS analysis was carried out using a model 6540 Accurate-Mass Q-TOF LC/MS (Agilent Technologies, Palo Alto, CA, USA), equipped with an electrospray ion source with Agilent Jet Stream technology operating in positive ionization mode. Data acquisition was performed in full scan mode in the mass range of 100–800 m/z. The source parameters were set as follows: gas temperature at 320°C, gas flow at 8 L.min$^{-1}$, nebulizer pressure at 30 psi, sheath gas temperature at 375°C, sheath gas flow at 12 L.min$^{-1}$, VCap at 3750 V, nozzle voltage at 500 V, and fragmentor voltage at 150 V. For continuous mass calibration, the following reference ions were used: purine 121.0508 $[M + H]^+$ and HP-921 = hexakis (1H, 1H, 3H-tetrafluoropropoxy) phosphazine 922.0097 $[M + H]^+$. Data acquisition and elaboration was carried out using the software MassHunter Acquisition and Qualitative Analysis version B.04.00 (Agilent Technologies) in accordance with (Gottardo et al., 2014). HPLC analysis was carried out using a Shimadzu LC-2030 Plus Prominence-i (Japan) system equipped with a Shimadzu standard UV-detector. Chromatographic separation was carried out by gradient elution using a TSKgel Protein C4-300 column (15×4.6 mm, Tosoh Bioscience). The composition of mobile phase was as follows: 0.1% (v/v) TFA (solvent A), and 0.1% (v/v) TFA in acetonitrile (solvent B). The eluents A and B were filtered through 0.2 µm pore size filter. The injection volume for all samples was 5 µL and the flow rate was 0.5 mL.min$^{-1}$ whereas the separation was obtained following gradient program: 0–4 min 20% solvent B, 4–19 min from 20% to 100% solvent B, 19–24 min 100% solvent B, 24–28 min 20% solvent B (re-equilibration step). The column and UV-detector were both maintained at 40°C throughout analysis. All the data acquired were processed by Shimadzu LabSolutions control software.

**Method S7. Bioinformatic analyses.**

Protein sequences of OSOXs (FASTA format) were obtained from UniProt database (https://www.uniprot.org) and used for the multiple amino acid alignment by using the online Align tool (http://www.uniprot.org/align) and Clustal Omega program (https://www.ebi.ac.uk/jdispatcher/msa/clustalo).

**Table S1.** Primers used in this work.

| GENE | ATG CODE | FORWARD PRIMER (5’->3’) | REVERSE PRIMER (5’->3’) |
|---|---|---|---|
| ***UBQ5*** | AT3G62250 | GTTAAGCTCGCTGTTCTTCAGT | TCAAGCTTCAACTCCTTCTTTC |
| ***FOX1*** | AT1G26380 | AGGTTCTCGAACCCTAACAACA | GCACAGACGACACGTAAGAAAG |
| ***FRK1*** | AT5G51830 | TTAAACTCGACGATGCAACA | GATGGAAGTTTTCCCGTTTT |
| ***CYP81F2*** | AT5G57220 | GTGAAAGCACTAGGCGAAGC | ATCCGTTCCAGCTAGCATCA |
| ***WRK33*** | AT2G38470 | GGCTCATCGATTGTCAGCAG | CCTTAACGACTTTCTGGCCG |
